# Aggregation of Transthyretin by Fluid Agitation

**DOI:** 10.1101/2024.11.08.622726

**Authors:** Irina Ritsch, H. Jane Dyson, Peter E. Wright

## Abstract

The transthyretin (TTR) tetramer, assembled as a dimer of dimers, transports thyroxine and retinol binding protein in blood plasma and cerebrospinal fluid. Aggregation of wild type or pathogenic variant TTR leads to transthyretin amyloidosis (ATTR), which is associated with neurodegenerative and cardiac disease. The trigger for TTR aggregation under physiological conditions is unknown. The tetramer is extremely stable at neutral pH, but aggregation via tetramer dissociation and monomer misfolding can be induced *in vitro* by lowering the pH. To elucidate factors that may cause TTR aggregation at neutral pH, we examined the effect of shear forces such as arise from fluid flow in the vascular system. Fluid shear forces were generated by rapidly stirring TTR solutions in conical microcentrifuge tubes. Under agitation, TTR formed β-rich aggregates and fibrils at a rate that was dependent upon protein concentration. The lag time before the onset of agitation-induced aggregation increases as the total TTR concentration is increased, consistent with a mechanism in which the tetramer first dissociates to form monomer that either partially unfolds to enter the aggregation pathway or reassociates to form tetramer. NMR spectra recorded at various time points during the lag phase revealed growth of an aggregation-prone intermediate trapped as a dynamically perturbed tetramer. Enhanced conformational fluctuations in the weak dimer-dimer interface suggests loosening of critical inter-subunit contacts which likely destabilizes the agitated tetramer and predisposes it towards dissociation. These studies provide new insights into the mechanism of aggregation of wild type human TTR under near physiological conditions.

## Introduction

Transthyretin (TTR) is a protein component of blood plasma and cerebrospinal fluid that functions in the transport of thyroid hormones and retinol-binding protein (1–5). Misfolding and aggregation results in neurological and cardiac TTR amyloidosis (ATTR) (6), which may be associated either with inherited amyloidogenic TTR variants, or with deposition of wild-type (WT) TTR fibrils in the heart (ATTRwt). ATTRwt is a disease of aging that affects up to 25% of the population over the age of 80 (7).

Transthyretin forms a stable tetramer in solution (8, 9) (Figure 1); aggregation and amyloid formation are thought to involve dissociation of the tetramer and misfolding of the resulting monomeric forms. Recently-reported high resolution structures of amyloid fibrils isolated from tissues of ATTR patients have revealed a complex cross-β-sheet topology, both for spontaneous ATTRwt (10), and hereditary ATTR (11–14). Each of these structures requires unfolding and remodeling of the native TTR structure. The overall similarity of the fibril core across different mutations and tissues is striking, and likely indicates a conserved pathway of formation (10, 14).

**Figure 1.**
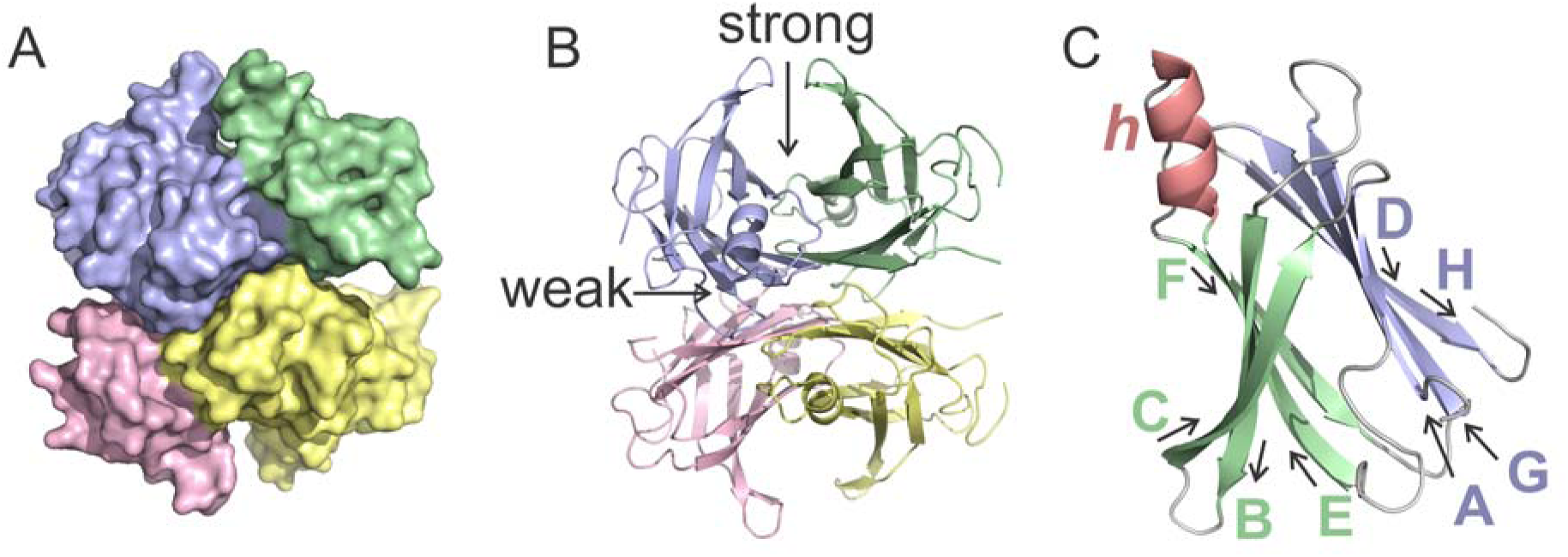
A. space-filling model of the structure of native TTR tetramer colored by subunit; B. ribbon model of the tetramer showing strong and weak dimer interfaces. C. ribbon model of one protomer with annotation of the β-strands (A-H), and the EF helix *h* (pdb: 1TTA) (25).

Under physiological conditions *in vitro*, neither the WT tetramer nor an engineered monomer (M-TTR) aggregate spontaneously, even over long time periods (15). Aggregation studies *in vitro* typically use acidic conditions (pH 4.4) to accelerate both tetramer dissociation and monomer misfolding (16). The monomer is not intrinsically amyloidogenic and must undergo partial unfolding to enter the aggregation pathway (15, 17). Extensive study of the acid-induced aggregation pathway of TTR led to the development of small molecule kinetic stabilizer drugs that suppress tetramer dissociation and thereby block the first step in the aggregation cascade (18).

The factors that trigger TTR aggregation *in vivo* remain unknown. Amongst postulated causes are the acidic environment of late endosomes and lysosomes, (19) age-related impairment of proteostasis networks (20, 21), oxidative damage (22), and proteolytic cleavage enabled by shear and interfacial forces associated with fluid flow (23, 24).

Elucidation of the mechanism of transthyretin aggregation and amyloid formation under physiological conditions is of major importance. In the present work, we ask whether shear and interfacial forces from fluid flow, which are ubiquitous in the vascular system (26), could be operational in the genesis of TTR amyloid. We describe an experimental approach to apply stress to the native protein *in vitro* by rapid stirring (agitation) as a mimic of naturally turbulent fluid flows in the bloodstream (27, 28). Agitation has previously been reported as an accelerator of aggregation kinetics for acid denatured TTR (29) or in the presence of proteases (23, 24, 30, 31). Here we focus on isolating the effect of agitation on otherwise unperturbed full length TTR, under close to physiological conditions, at neutral pH. The most extreme blood flow conditions are found in the heart and aorta (27, 28, 32), and our results are expected to be relevant primarily to cardiac amyloidosis.

## Materials and Methods

### Preparation of Proteins

Plasmids containing the genes for human TTR and the monomeric F87E variant were used as previously described (16, 33). Full details of protein preparation and purification are given in Supporting Information. After tag cleavage, all TTR constructs retained an additional Gly that precedes the native N-terminal Gly residue. Reducing agent (1 mM tris(2-carboxyethyl)phosphine, TCEP) was present during the purification and in the final stock solutions (∼1 mM TTR) in agitation buffer (also used for storage) (10 mM potassium phosphate, pH 7.0, 100 mM KCl, 1 mM EDTA); these solutions were aliquoted (∼200 μl), snap frozen in liquid nitrogen and stored at −80°C until usage (no further freeze-thaw cycles). High purity (>95%) after the second Ni-affinity step was confirmed for all protein samples by SDS-PAGE, reverse phase HPLC, and MALDI mass spectrometry. Unless stated otherwise, protein concentrations are given as protomer concentration determined by UV extinction at 280 nm with specific molar extinction coefficient ε = 18450 M^-1^ cm^-1^ calculated with Protparam (34).

### Acid aggregation assay

Aggregation assays were performed by diluting protein stock solutions in agitation buffer to the desired initial concentration (*c*_0_) in 50 mM sodium acetate buffer, pH 4.4, 100 mM KCl. For each time point, three independent sealed tubes were incubated at 37°C without shaking

### Agitation assay

The agitation assay was performed in agitation buffer. Stock TTR solutions were diluted in agitation buffer to the desired concentration and left to equilibrate for at least 1h before starting agitation. Protein samples were individually agitated by rapid stirring (1200 rpm) with a single rectangular, Teflon coated, magnetic stirring flea (Fisher Scientific, 7 x 2 x 2 mm) in 1.5 ml conical microcentrifuge tubes (Costar) on linked modules of a multi-plate magnetic stirrer (Cole-Parmer UX-84003-80). Agitation was performed at ambient temperature and to minimize sample heating and evaporation, the microcentrifuge tubes were sealed with parafilm and immersed in a water bath. Three independent samples (n=3) were agitated without interruption in separate containers for each time point. To avoid errors from small variations in protein and buffer batches, comparative agitation screens were run in a concerted manner in the following series:

(1) concentration dependence screen with 5, 10, 20, 50, 100 μM TTR and 5, 10, 20 μM F87E for scattering and fluorescence analysis (250 μl per tube)
(2) supernatant Trp fluorescence screen with 0.5, 1, 5, 10, 50, 100 μM TTR (250 μl per tube)

To assess the role of the air-water interface (AWI) in agitation-induced aggregation of TTR, we modified microcentrifuge tubes to eliminate the AWI. Each tube was cut at the 750 μl line and filled with 5 μM TTR in agitation buffer. A second microcentrifuge tube was then inserted to squeeze out excess liquid and eliminate the air gap and the assembly was sealed with parafilm. Three independent samples (∼ 700 μl) were stirred at 1200 rpm at ambient temperature for 72 hours, in parallel with two reference samples (800 μl) in unmodified microcentrifuge tubes with an air-water interface. The extent of aggregation was assayed using light scattering at 600 nm and thioflavin T (ThT) fluorescence (see below).

### Light extinction (scattering and soluble TTR assays)

The aggregation of TTR was estimated in two ways, ‘total’ analysis, an estimate of the extent of precipitation by light scattering at 600 nm (*A*_600nm_), and ‘supernatant’ analysis, a measurement of the remaining soluble protein after centrifuging the sample, by absorbance at 280 nm (*A*_280nm_). Light extinction was measured on an Infinite® 200 PRO plate reader (Tecan).

For ‘total’ sample analysis we used 80 μl sample per well in transparent, unlidded, 96 well plates (Corning^TM^, non-treated, flat bottom). For ‘supernatant’ protein analysis, samples were centrifuged for 30 min at 17k x g to remove large aggregates and assayed using 100 μl sample per well in UV-transparent, unlidded, 96 well plates (Part No. 8404, Thermo, UV Flat Bottom Microtitor®). All plate reader data were baseline-corrected using reference data from buffer-only wells.

### ThT fluorescence measurements

After measurement of the scattering intensity, the samples were immediately re-used for the ThT assay. ThT was added from stock solution (100 mM in phosphate buffer) to a final ThT concentration of 100 μM and left to incubate in the dark without shaking for 10 min. ThT fluorescence was measured by excitation at wavelength λ_ex_ = 440 nm and emission at wavelength λ_em_ = 480 nm. The increase in fluorescence intensity was determined relative to the intensity of free ThT in agitation buffer (ThT gain = *I*_ThT_*/I*_ThT,free_).

### Tryptophan fluorescence

Tryptophan fluorescence (screen 2 above) was measured after agitation, with ‘supernatant’ fractions (centrifuged 30 min at 17k x *g*) from the TTR concentration dependence screen (screen 1 above). Trp fluorescence was measured on a Horiba Fluorolog spectrometer at 298 K. To avoid concentration-dependent differences in fluorescence quenching, each sample was diluted to 0.25 μM TTR (in agitation buffer) as a constant volume ratio dilution for each *c*_0_ (i.e. not corrected for loss of soluble TTR due to aggregation). Fluorescence emission spectra maximum (λ_em_ = 345 nm) *I*_w_(*t*)was extracted after baseline subtraction of a buffer-only (λ_em_ = 305-500 nm, λ_ex_ = 280 nm, step size 1 nm) were recorded and the intensity at the emission measurement.

### Negative stain transmission electron microscopy

A highly ThT positive, late-stage sample (43 day agitation, *c*_0_ = 5 μM) was gently sonicated in a water bath for 1h to reduce the size of very large aggregates, and then gently centrifuged for 5 min at 2k x g. The supernatant (3 μl) was loaded onto a glow-discharge activated carbon-coated copper grid (1 min incubation before blotting). To suppress stain crystallization artifacts from phosphate buffer, blotted grids were washed with two drops of double distilled water (3 μl each, instant blotting). Samples were stained with two cycles of uranyl formate buffer (10s, 30s). Images were collected using the Leginon software (35, 36) on an FEI Talos electron microscopes at 73,000x magnification (pixel size 1.981 Å).

### Fitting of aggregation time-course data

Fitting of aggregation time courses (induced by acid or agitation) was performed in MATLAB using custom scripts. The increase in intensity *I*(*t*) for *A*_600nm_ and ThT fluorescence as a function of stress application time *t* were fitted with a three-parameter sigmoidal model (Equation 1):

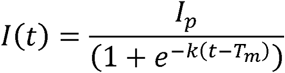

with plateau intensity *I*_p_, exponential phase rate *k*, and midpoint time *T*_m_. The fit model assumes that any offset intensity *I* (t=0 h) = *I*_0_ at the beginning of the assay is negligible, which was experimentally confirmed (*I*_0_ ≈ 0 ≪ *I*_p_).

The decrease in supernatant Trp fluorescence intensity as a function of *t* was fitted with a modified sigmoidal function (Equation 2)

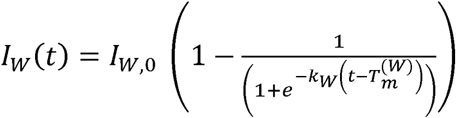

where *I*_*w*,0_ is the initial intensity and *k*_W_ and *T*_m_^(W)^ are the exponential rate constant and the mid-point time of the fluorescence signal decay, respectively. The model assumes that the fluorescence intensity converges to zero once aggregation is complete *I* (t→∞) ≈ 0.

### NMR analysis of supernatant

Solution state NMR spectroscopy was performed with agitated samples of uniformly ^15^N labeled wild-type TTR in agitation buffer. Despite the high molecular weight of the native tetramer (∼56 kDa) protein deuteration was not used since it is known to affect intrinsic TTR stability (25). In addition to an unperturbed reference sample (time *t*_0_), samples for the NMR experiments were subjected to 4 – 21 days of continuous agitation (1200 rpm) at ambient temperature. ThT gain and *A*_600nm_ were determined as end-point measurements in a 96-well plate as described above, using a small aliquot of ‘total’ samples after 1:10 dilution in agitation buffer (to reduce the sample volume required at high concentration). As a control to confirm that trends in the samples hold after dilution, we also measured *A*_330nm_ on the nanodrop in UV/Vis mode for each sample prior to dilution. To prevent interference from large aggregates with the long-term stability of shimming they were removed by centrifugation (30 min at 17k x g). Final samples after addition of 10% D_2_O for spectrometer locking were transferred to Shigemi tubes, and ^1^H-^15^N HSQC spectra were acquired at 298K on a Bruker Avance 700 spectrometer with a ^1^H/^19^F– ^13^C/^15^N TCI cryoprobe and shielded z-gradient coil. Spectra were processed using NMRpipe (37), and assignments were transferred from those of the wild-type TTR tetramer (BMRB 27514) (38) after minor chemical shift adjustments to account for differences in buffer composition. Cross-peak chemical shifts, linewidths Γ, and intensities *I* were extracted with a custom MATLAB script using the reference peaklist and allowing small variations in peak position (<0.05 ppm in ^1^H, < 0.2 ppm in ^15^N) and linewidth of a 2D Gaussian peak. The changes in HSQC cross peak intensity at each time point relative to the non-agitated reference sample were monitored using intensity ratios, I(t)/I_0_. For each of the independent samples, the intensities were normalized using the average intensity of residues 3-6 in the disordered N-terminal region.

## Results

### Agitation by rapid stirring causes aggregation of human TTR at neutral pH

To measure the effects of agitation on the aggregation of TTR we utilized an in-house setup where Eppendorf tubes with stir bars were employed with a precision stirring motor to give reproducible room-temperature agitation. The time course of aggregation (of the order of days or weeks) was then assayed using turbidity and residual protein concentration measurements and light microscopy.

Since the standard method of studying TTR aggregation is by the application of acidic conditions, we first made a comparison of the effects of low pH (4.4) and of agitation on a human TTR sample (initial protomer concentration *c*_0_ = 5 μM). Both acid pH (under quiescent conditions) and agitation (at 1200 rpm in agitation buffer in individual conical microcentrifuge tubes) resulted in sample turbidity and the appearance of aggregates observable by light microscopy after three days. Plots of turbidity (measured as light extinction at 600 nm; *A*_600nm_) as a function of time are shown in Figure 2A, B.

**Figure 2.**
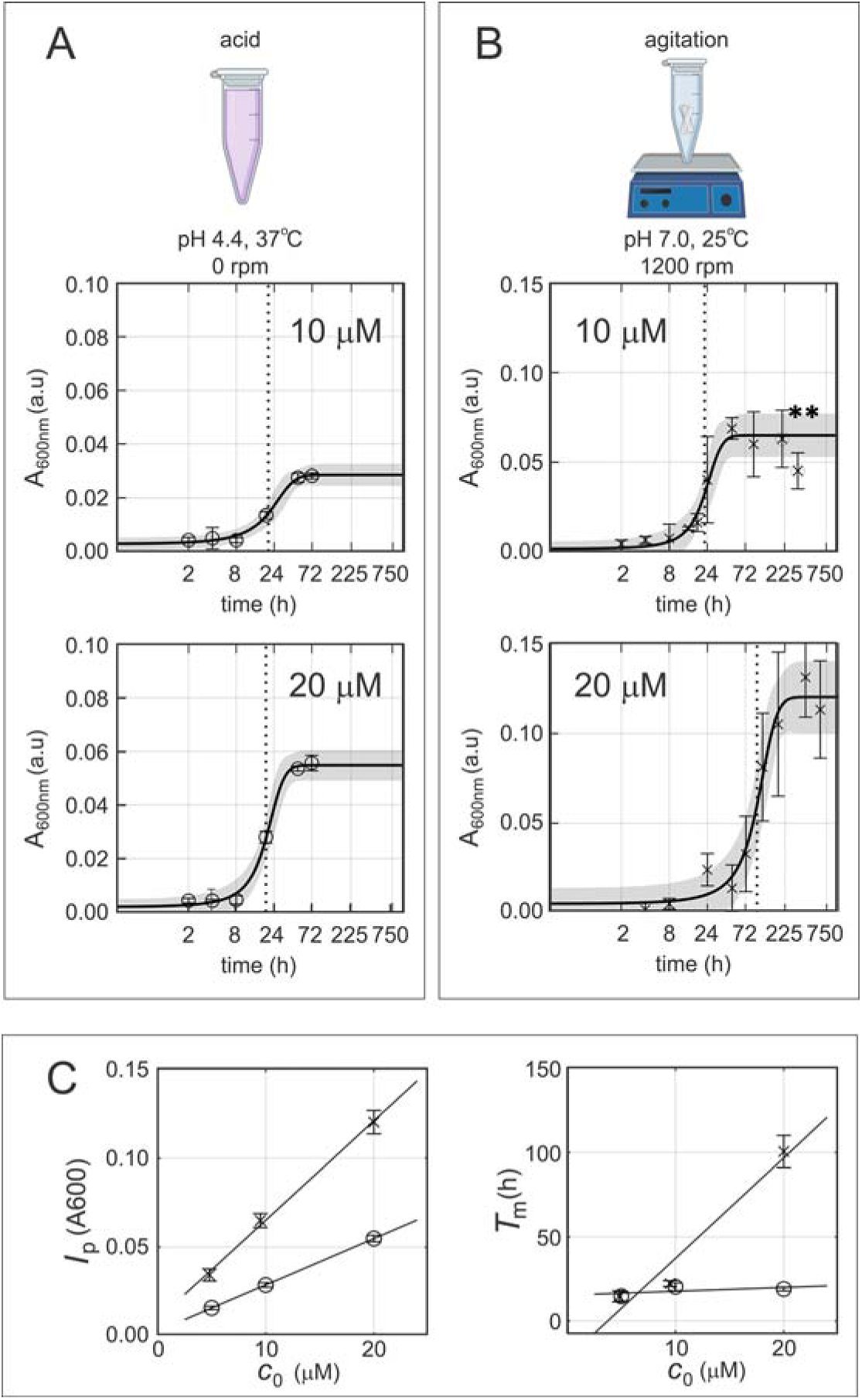
Comparison of the time course of TTR aggregation by A. acid and B. agitation, measured by light scattering at 600 nm (*A*_600nm_) for *c*_0_ =10 μM, and 20 μM. Error bars represent the standard deviation of three independent samples per individual time point. Solid black line is a sigmoidal fit with Equation 1, with 90% confidence intervals shaded grey. Dotted vertical lines mark *T*_m_. Asterisks mark time-points that showed loss of *A*_600nm_ after a peak value. C. Concentration dependence of the fit parameters of *A*_600nm_ assays for *c*_0_ = 5-20 μM for agitation (x) and acid (o) conditions; solid lines show linear regressions; the slope of the *I*_p_ vs *c*_0_ regression is 1.06 ± 0.05 μM^-1^ for agitation, and 0.6 ± 0.03 μM^-1^ for acid.

### Concentration dependence of TTR aggregation

Turbidity measurements were made for TTR solutions over a range of starting concentrations (*c*_0_ = 5-100 μM protomer) spanning the physiologically relevant range in blood (∼7-29 μM) (39). The kinetic profiles for acid-induced and agitation-induced aggregation (shown for two concentrations in Figure 2 and for all concentrations in Figure S1) show an initial lag phase, followed by an exponential phase before converging to a plateau value. Aggregation parameters (midpoint time *T*_m._, plateau scattering intensity *I*_p_, and exponential phase aggregation rate *k*) were extracted by fitting the aggregation profiles to a three-parameter model given in Equation 1 (solid lines in Figure 2A, B and Figure S1).The fitted values under all conditions are shown in Table 1.

**Table 1.** Fitted parameters for TTR aggregation (*A_600nm_* time courses) by acid or agitation.

| $c_0$ [$\mu\text{M}$] | $k$ ($A_{600\text{nm}}$)<br>[1/h] | $I_p$ ($A_{600\text{nm}}$)<br>[a.u.] | $T_m$ ($A_{600\text{nm}}$)<br>[h] | $f_M$ | $R_{\text{adj}}$<br>($A_{600\text{nm}}$) |
| --- | --- | --- | --- | --- | --- |
| ACID |  |  |  |  |  |
| 5 | $0.09 \pm 0.03$ | $0.017 \pm 0.003$ | $18 \pm 3$ | n.d. | 0.99 |
| 10 | $0.11 \pm 0.03$ | $0.029 \pm 0.002$ | $20 \pm 2$ | n.d. | 0.99 |
| 20 | $0.18 \pm 0.04$ | $0.055 \pm 0.003$ | $19 \pm 1$ | n.d. | 0.99 |
| AGITATION |  |  |  |  |  |
| 5 | $0.2 \pm 0.1$ | $0.03 \pm 0.01$ | $15 \pm 3$ | 0.0065 | 0.79 |
| 10 | $0.15 \pm 0.05$ | $0.07 \pm 0.01$ | $24 \pm 2$ | 0.0039 | 0.98 |
| 20 | $0.03 \pm 0.01$ | $0.13 \pm 0.01$ | $106 \pm 9$ | 0.0023 | 0.97 |
| 50 | $0.01 \pm 0.01$ | $0.3 \pm 0.3$ | $200 \pm 140$ | 0.0012 | 0.77 |
| 100 | $0.01 \pm 0.01$ | $0.6 \pm 2$ | $300 \pm 400$ | 0.00069 | 0.88 |

Aggregation of TTR by acid and agitation at different initial protomer concentrations ( $c_0$ ), observed by light scattering $A_{600\text{nm}}$ and fitted to Equation 1. Errors are shown at 90% fit confidence interval. $k$ : exponential phase aggregation rate; $I_p$ : plateau scattering intensity; $T_M$ : midpoint time; $f_M$ : fraction of monomer; $R_{\text{adj}}$ : adjusted goodness of fit.

For both acid and agitation-induced aggregation, the *I*_p_ increases linearly with concentration (Figure 2C), but is consistently higher under agitation. The linearity confirms that within each stress condition, the samples converge to a similar aggregate composition, and the differences in slope and absolute values indicate that aggregate morphology and scattering properties likely differ between the aggregation protocols. A more obvious difference is observed between the two protocols in the concentration dependence of *T*_m_, which was close to independent of *c*_0_ with acid denaturation but showed a substantial increase at higher *c*_0_ with agitation (Figure 2B). The same trends are seen under agitation at higher TTR concentrations (Figure S1), but a well-defined plateau phase was not reached at these higher concentrations, and we thus refrain from detailed interpretation of the fitted parameters in this range.

The observed increase in lag time with increasing TTR concentration is unusual and is the opposite of that for a typical nucleated aggregation process (40, 41). The inverse concentration dependence of *T*_m_ is consistent with a mechanism in which the aggregation-prone species is the monomer. Under agitation, the monomer can either enter the aggregation pathway or reassociate to form the native TTR tetramer which acts as an off-pathway oligomer that sequesters the TTR protomers in a non-aggregating form (42–44). Using the tetramer dissociation constant for human TTR (9 x 10^-25^ M^3^) (39), we calculated the fraction of monomer at each TTR concentration (*f_M_*= [M]^4^/c_0_, where c_0_ = 4[T]+[M] is the total TTR concentration) as described previously (45). The lag time *T*_m_ is inversely correlated with the fraction of monomer *f_M_*, which decreases as the concentration c_0_ increases (Table 1). A plot (46) of log *T*_m_ vs log *f_M_* is linear with slope −1.4 (Figure S2).

To gain further insights into the mechanism of agitation-induced aggregation, we repeated the assay with an engineered monomeric variant (F87E) of TTR (33). Like TTR, F87E aggregates under agitation but not under quiescent conditions in the time frames and concentration ranges in our experiments. Agitation-induced aggregation of wild type TTR and F87E at multiple concentrations was monitored using the increase in thioflavin T fluorescence. The time-course of agitation-induced ThT fluorescence gain at two concentrations is shown in Figure 3A, with fitted parameters in Table S1. At *c*_0_ = 5 and 10 μM, the lag time for agitation-induced aggregation of F87E was similar to that of the WT tetramer and increased only slightly at higher concentrations (*c*_0_ = 20 μM, see Figure 3B), confirming that the pronounced concentration-dependent increase in lag time for aggregation of WT TTR is suppressed in a variant that cannot form a stable tetramer. The modest concentration-dependent increase in *T*_m_ for F87E is due to a propensity to self-associate to form an off-pathway oligomer at the relatively high KCl and phosphate concentration of the agitation buffer.

**Figure 3.**
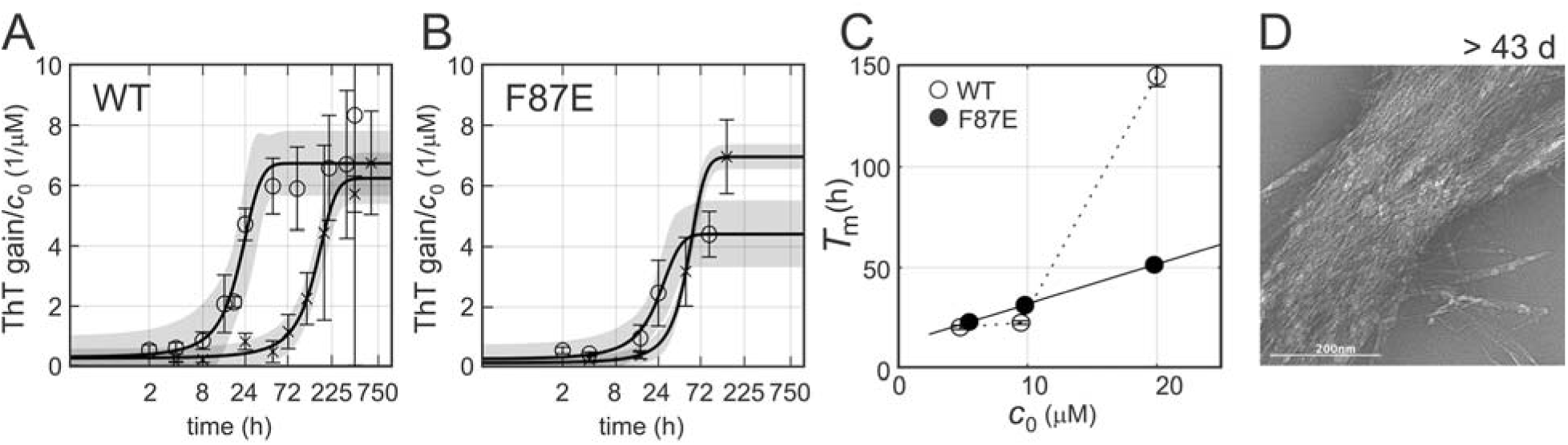
A. Normalized time course of ThT gain = *I*_ThT_ */I*_ThT,ref_ for two initial protomer concentrations, *c*_0_ = 5 μM (o), or 50 μM (x) and fits with Equation 1 (90% confidence intervals shaded grey) for WT TTR and B. for the F87E monomeric variant. C. fitted lag times *T*_m_ for WT and F87E ThT gain at three values of *c*_0_; D. negative stain TEM of WT fibrils (*c*_0_ 5uM) after 43 d agitation.

### Aggregate morphology

To examine the morphology of the late stage, highly ThT-positive aggregates we performed negative stain transmission electron microscopy (TEM). The final stage aggregates (43 d) were clearly fibrillar, with individual protein fibers arranged in large, intertwined bundles (Figure 3D). Virtually no residual particles consistent with the size of the TTR tetramer (∼55 kDa) could be detected.

### Role of the air-water interface

Agitation-induced aggregation can be potentiated by fluctuations at the hydrophobic air-water interface (AWI) or, in the absence of an air interface, by turbulent shearing forces at the interface between the liquid and the surface of the container (47, 48). To assess the role of the AWI in agitation-induced TTR aggregation we stirred a solution of WT TTR (*c_0_* = 5 μM in agitation buffer) in Eppendorf tubes at a rate of 1200 rpm for 72 hours in the presence or absence of an air gap. Agitation yielded comparable amounts of ThT positive aggregates (*A_600_* = 0.021, I_ThT_/I_ThT0_ = 4 ± 1 with an air gap and *A_600_* = 0.023, I_ThT_/I_ThT0_ = 8 ± 3 with no air gap), suggesting that the AWI is not strictly required for agitation-induced TTR aggregation.

### Monitoring TTR aggregation using tryptophan fluorescence spectroscopy

To confirm that the observed aggregates contain TTR we measured the loss of supernatant (soluble) Trp fluorescence emission intensity I_w,sol,_ after removal of large insoluble aggregates by centrifugation. Aliquots were taken from the sample solution at various time points during agitation and Trp fluorescence spectra were measured after dilution. Results for two concentrations are shown in Figure 4A, and for all concentrations in Figure S3. As expected, I_w,sol_ decreased as a function of agitation time and, consistent with the scattering assay, only a short lag phase was observed for *c*_0_ = 5 μM (Figure 4A). Given the high sensitivity of the Trp fluorescence assay, we could extend the measurements to concentrations as low as *c*_0_ = 0.5 μM. No discernable lag phase was observed below *c*_0_ = 5 μM (Figure S3). In contrast, we observed an extended lag phase at *c*_0_ = 50 μM, during which I_w,sol,_ slightly increased (Figure 4A, B). The initial intensity gain likely arises from either a population of small oligomers that are not removed by centrifugation or from changes in the conformation or dynamics of the tetramer that transiently relieve the intrinsic fluorescence quenching of Trp79 in the native state (16, 49). We fitted a modified sigmoidal model (Equation 2) to the Trp fluorescence data to determine the initial intensity I_W,0_, midpoint time T_m(W)_, and exponential phase rate k_W_ (Table S2). As observed with *A*_600nm_ assays, T_m_ increased at higher *c*_0_. Other than changes in Trp fluorescence intensity, no substantial spectral changes were observed that could reveal the conversion of folded to fully or partially unfolded protomers (Figure 4B).

**Figure 4.**
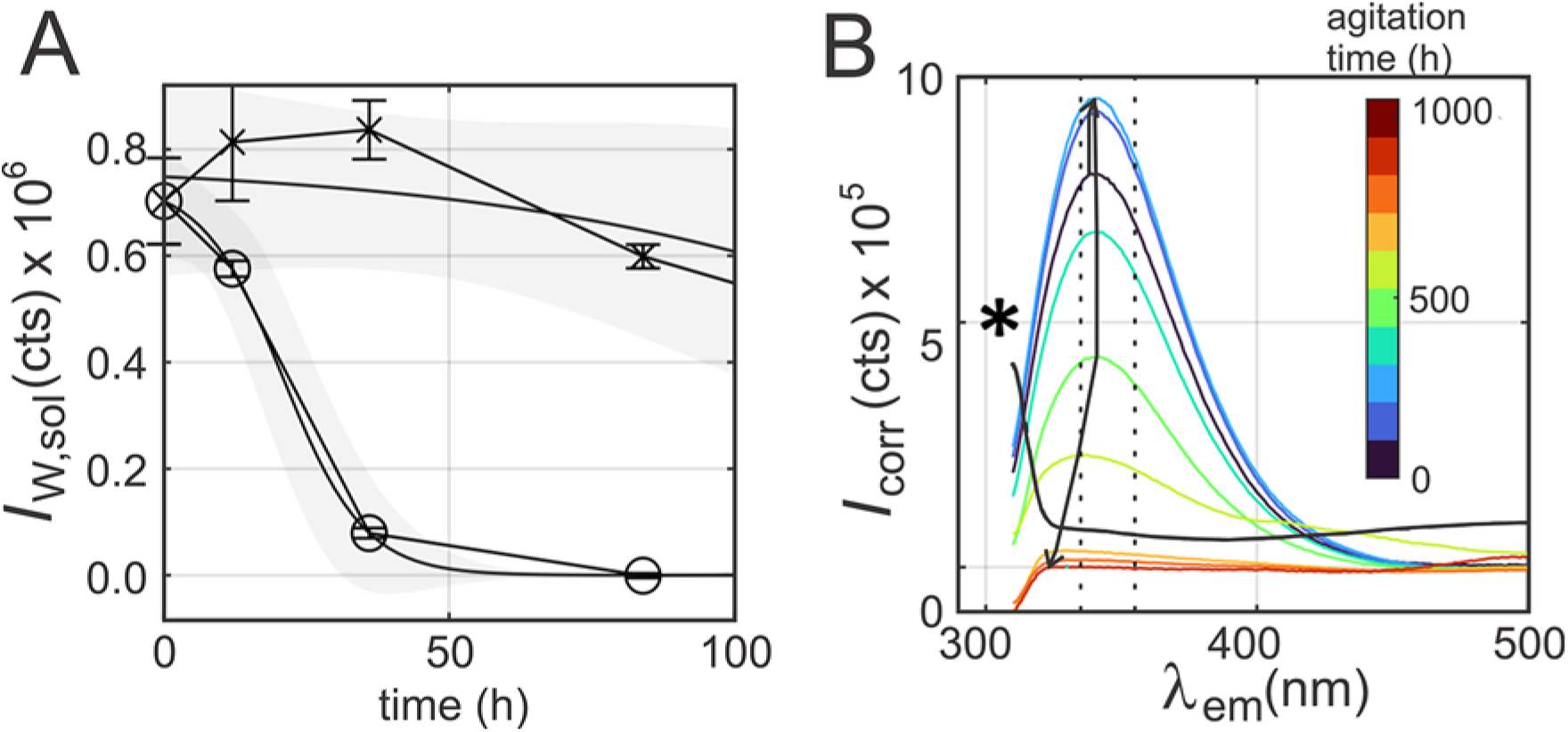
(A) Supernatant (‘soluble’) native Trp fluorescence intensity, *I*_W,sol_, at 345 nm (ex 280 nm) after large aggregate removal by centrifugation. *c*_0_ = 5 μM (o) and 50 μM (x), and fits with Equation 2 (90% confidence intervals shaded grey); (B) full fluorescence emission spectra for 50 μM time-course shown in (A), and blank (dash-dotted), asterisk marks position of solvent Raman peak in blank. The arrow marks the transient *I*_W,sol_ increase, and vertical lines mark positions of folded (em 335 nm), and unfolded (em 355 nm) spectral maxima under urea denaturation.

### NMR analysis of TTR in the lag phase

To obtain insights into the structural and dynamic changes in the TTR tetramer induced by agitation, we acquired ^1^H-^15^N HSQC spectra of the supernatant at various time points during agitation by a magnetic stir bar in conical centrifuge tubes. Since deuteration destabilizes TTR (25), we used a high concentration of non-deuterated TTR (*c*_0_ = 350 μM) to ensure a good signal-to-noise ratio for both weak and broad cross peaks. The kinetics of aggregation are slow at this high concentration and agitation was thus performed for relatively long periods (4 to 21 days). All time points sampled are well within the lag time which, by extrapolation of a linear plot of *T_m_* versus *c*_0_ (Figure S4A), is estimated to be ∼45 days at 350 μM concentration. From the light scattering amplitude at the 21-day endpoint of this experiment (*A*_600nm_ = 0.3, Table S3) and by extrapolation of the observed linear relationship between *I*_p_ *(A*_600nm_) and concentration *c*_0_ (Figure S4B), we estimate that the extent of aggregation is ∼14%, well within the lag phase.

The species formed during the lag phase are of particular interest, promising to provide insights into the structural and dynamic changes in TTR that are induced by agitation and drive entry into the aggregation cascade. A superposition of the ^1^H-^15^N HSQC spectrum of the supernatant after 21 days of agitation and that of an unperturbed reference sample is shown in Figure 5A. The cross peak positions are unchanged, showing that the residual soluble protein in the agitated sample adopts a native-like tetrameric state.

**Figure 5.**
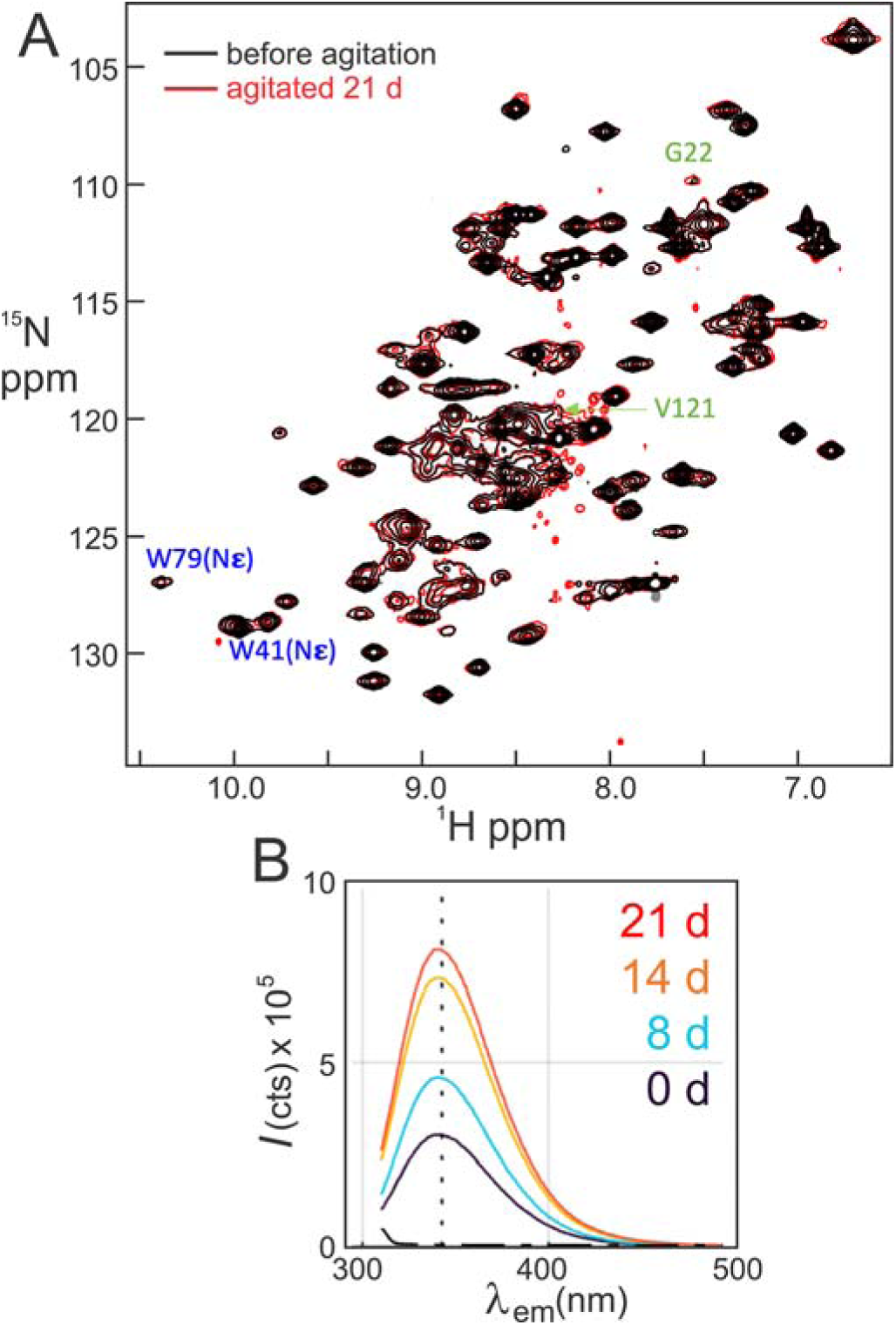
A. Superposition of ^1^H-^15^N HSQC NMR spectra (298 K, 700 MHz) of supernatant TTR fraction before (black) and after 21 d agitation (red); W41, and W79 side chain peaks are marked blue. Resonances of G22 and V121, which show intensity changes are labeled in green. B. *I*_W,sol_ of the supernatant TTR NMR samples including those shown in part A at agitation times: t = 0 reference (black), 8 days (blue), 14 days (orange) and 21 days (red).

While the chemical shifts are unchanged by agitation, the intensities of many cross peaks change substantially relative to the reference spectrum. Histograms summarizing the intensity changes at various agitation time points are shown in Figures 6 and S5. To correct for potential differences in spectrometer sensitivity or shimming, intensities at each time point were normalized using the average intensity of the cross peaks of residues 3-6, which remain disordered under all conditions.

**Figure 6.**
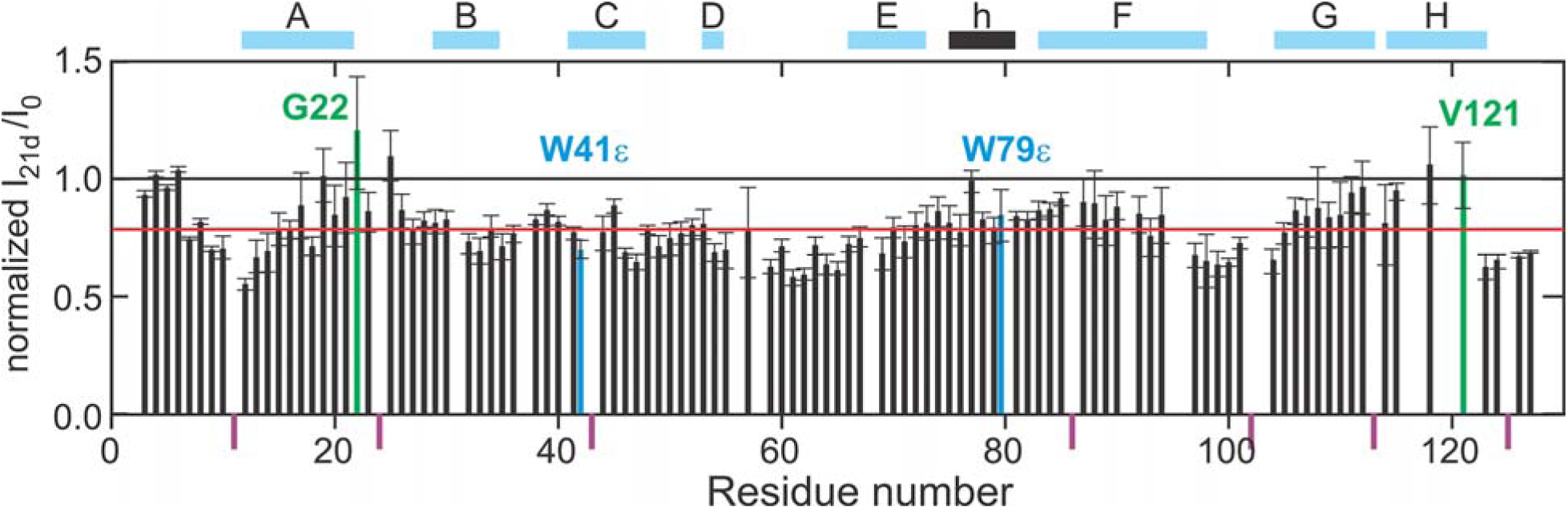
Ratio of the cross peak intensity after 21 days agitation I_21d_ to the intensity before agitation I_0_ in the ^1^H-^15^N HSQC spectra of supernatant TTR after agitation, normalized to the average intensity ratio of the cross peaks of residues 3-6. The black solid line indicates no change, red solid line indicates average ratio (0.8). Pro residues are marked by purple bars below the x-axis, data for tryptophan side-chain NεH resonances are shown in blue and for two interface residues discussed in the text in green. The location of secondary structure elements is indicated at the top of the figure: A-H β-strands; h EF helix.

A general decrease in cross peak intensity was observed after agitation, with greater intensity loss at longer agitation times (Figures 6, S5). However, not all cross peaks lose intensity at a uniform rate; cross peaks for several residues lose intensity more rapidly than average while, unexpectedly, the intensities of some cross peaks remain unchanged or even increase with increasing agitation time. The observed differential cross peak intensity changes must reflect changes in local dynamics, since the chemical shifts are not perturbed by agitation (which confirms that the ground state structure is the same as the native TTR tetramer). A gain in cross peak intensity after agitation implies decreased *R_2_*relaxation rates due to enhanced local dynamics in the ground state. Conversely, for cross peaks that decay faster than average, the intensity loss does not arise from loss of soluble protein but reflects increased *R_2_* relaxation rates (increased broadening), probably arising from exchange with a small population of a conformationally altered, tetrameric excited state. Residues that exhibit above-average intensity loss or gain (I/I_0_ < 0.72 or > 0.92, respectively) after 21 days of agitation are clustered in the TTR sequence and are shown in Figure 7 mapped to the structures of one TTR protomer (7A) and to the TTR tetramer (7B). Key sites of greater than average broadening/intensity loss map to the N-terminal region of the A-strand, strand D, the CD and DE and FG loops, and the C-terminal region of the H-strand. Residues that remain unchanged or gain in cross peak intensity are located at the C-terminal end of strand A, the AB loop, the GH loop, and V121 in strand H and all of these are clustered in the weak dimer interface (Figure 7B). One striking example is residue G22 (labeled in green in Figures 5A and 6) which is located at the weak dimer interface in close contact with V121 and Y114 of neighboring subunits (Figure 7C). The increased amide cross peak intensity of G22 suggests enhanced dynamics in the weak dimer interface associated with agitation, an effect that is also observed for the V121 cross peak.

**Figure 7.**
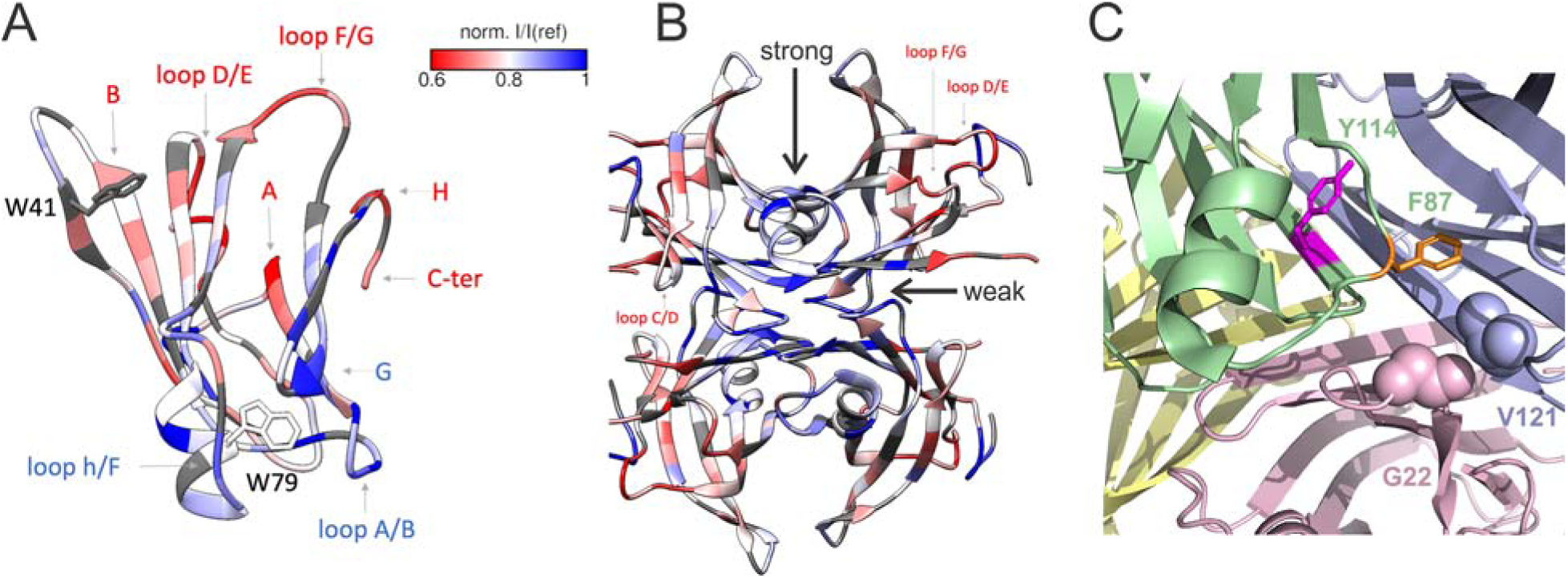
A. Intensity ratios from Figure 6, mapped onto one protomer of TTR; tryptophan side chains are shown (pdb: 1TTA); B. mapping of intensity ratio onto the TTR tetramer; C. Interface region formed by A/B loop region with G22 (pink protomer), and V121 (blue protomer) on opposing sides of the weak dimer interface (residues shown as spheres), and nearby aromatic residues F87 and Y114 from the green protomer shown in orange and magenta.

The agitated NMR samples were diluted into fresh buffer for fluorescence measurements. In parallel with the HSQC cross peak intensity changes, the tryptophan fluorescence intensity of the NMR samples at 345 nm increases markedly with increasing agitation time (Figure 5B).

## Discussion

The factors that drive TTR aggregation *in vivo* remain poorly understood. The present experiments provide the first insights into mechanisms of aggregation of WT human TTR under near physiological conditions. At physiological pH and temperature, the WT human TTR tetramer is stable and highly resistant against aggregation. Previous studies of TTR aggregation mechanisms have, of necessity, been performed under non-physiological conditions (acid) to accelerate tetramer dissociation and partial unfolding of the monomer to promote entry into the aggregation pathway (15, 17, 50). While these studies have been informative, they provide no insights into how aggregation could be initiated *in vivo*. To obtain insights into factors that might potentially drive TTR aggregation under physiological conditions, we subjected the protein to shear forces associated with stirring in microcentrifuge tubes. Due to the conical shape of the tubes, the rotation of the magnetic stir bar is not smooth but undergoes random orientational oscillations, resulting in chaotic or turbulent fluid flow. At near physiological conditions of pH and salt concentration, both WT human TTR and the monomeric F87E variant aggregate to form β-rich fibrils when agitated by stirring at 1200 rpm.

For a typical nucleated aggregation process, the lag time decreases at higher protein concentrations (40, 41, 51). For wild type human TTR, however, the rate of agitation-induced aggregation at neutral pH decreases and the lag time increases as the total TTR concentration is increased (Table 1 and Figure 2). Such inverse concentration dependence has been attributed to assembly of off-pathway oligomeric species that compete with the on-pathway formation of oligomers and fibrils(42–44) Aggregation of TTR is consistent with a mechanism in which the tetramer must first dissociate to form monomer which, under the influence of agitation, either partially unfolds and enters the aggregation pathway or reassembles the tetramer. This mechanism is supported by the observation that the aggregation lag time *T*_m_ is inversely correlated with the mole fraction of monomer *f_M_* (Table 1). A plot of log *T*_m_ (measured from light scattering at 600 nm) vs log *f_M_* is linear with slope −1.4 (Figure S2), suggesting that the aggregation kinetics are likely dominated by secondary nucleation (46, 52). Interestingly, the slope is comparable to that of the corresponding log/log plot (slope = −1.5) for aggregation of monomeric M-TTR (F87M/L110M) at pH 4.4 (50), suggesting that partial unfolding of the monomer is required for entry into and progression down the aggregation pathway at both neutral and acid pH.

Aggregation of proteins under shear generally occurs at the air-water interface or the interface with the material of the containing vessel (47, 48, 53). In the case of TTR, aggregation does not appear to involve the air-water interface, since comparable levels of aggregate are observed in the presence and absence of an air gap. More likely, aggregation is driven by continuously regenerating interactions with the hydrophobic polypropylene surface of the microcentrifuge tubes (48). NMR relaxation dispersion measurements show that both monomeric M-TTR and F87E TTR at near neutral pH undergo conformational fluctuations that populate an amyloidogenic excited state in which the hydrogen bonding network of the DAGH β-sheet is perturbed (33, 54). We speculate that these fluctuations, which are amplified at acidic pH and drive aggregation, likely cause transient exposure of hydrophobic surfaces that interact with the surface of the microcentrifuge tube and promote agitation-induced aggregation. Alternatively, prolonged exposure to turbulent shearing forces may act directly to promote partial unfolding of the monomer or perturb the conformation or dynamics of the tetramer. Further work will be required to elucidate the detailed molecular mechanism by which shear stresses promote TTR aggregation and determine the role of surface interactions in the aggregation process.

Strong hydrodynamic shearing and extensional forces can in principle lead directly to protein deformation and destabilization, dependent on the size, sequence, and structural malleability of the protein (55, 56). Few studies on the effect of agitation and shear forces on the conformation and dynamics of globular proteins have been reported. Raman spectroscopy revealed reversible changes in the secondary structure of lysozyme under oscillatory flow conditions (57). More recently, high resolution NMR was used to identify shear-induced changes in structure and dynamics of folded and unfolded superoxide dismutase at single residue resolution (58) while hydrogen exchange mass spectrometry showed that agitation destabilizes local structure and increases the rate of local unfolding of certain monoclonal antibodies (59). In the particular case of transthyretin, shear and interfacial forces associated with sample shaking render the K48-T49 peptide bond susceptible to proteolytic cleavage (23).

To obtain insights into structural or dynamic changes in TTR induced by hydrodynamic shear stresses, we subjected ^15^N-labeled TTR samples to prolonged agitation and recorded ^1^H-^15^N HSQC spectra of the supernatant at various time points. Aggregation is very slow at the high concentration (350 μM) required for NMR, and the sample remained in the lag phase even after 21 days of continuous agitation. The NMR spectra reveal an aggregation-prone intermediate trapped as a dynamically perturbed tetramer. All cross peaks are at the chemical shifts of the WT tetramer, albeit with perturbed intensities. No cross peaks associated with monomer (60) were observed, implying that its equilibrium concentration is very low; aggregation of monomer is in competition with rapid reassociation to tetramer, which is expected to be on a seconds time scale at this high concentration (61).

Differential changes in cross peak intensity relative to the reference spectrum were observed at the earliest agitation time point (4 days) and increased in magnitude with increasing agitation time. The cross peak intensities of several residues in the AB and GH loops and V121 at the C-terminal end of strand H decayed significantly more slowly than average or even increased in intensity relative to the reference spectrum, indicating increased local dynamics induced by agitation. These residues form the weak dimer interface which, in the native tetramer, is stabilized by inter-subunit hydrophobic contacts and hydrogen bonds, between the A19 and G22 carbonyl oxygens and the amide protons of Y114 and V122, respectively, on neighboring protomers across the dimer interface. The AB loop is also stabilized within the protomer by intra-subunit hydrogen bonds to the GH region (L17 CO to L111 NH and A19 NH to L111 CO) and by a hydrogen bond between the D18 carboxyl and the side chain of Y78 (E helix). The observation of enhanced dynamics in the AB and GH loops following agitation suggests loosening of critical interactions in the weak dimer interface, which would likely destabilize the tetramer and predispose it towards dissociation. Mechanistically, dissociation of the TTR tetramer has been shown to occur by scission of the weak dimer interface followed by rapid dissociation of the resulting strong dimer (62, 63). The weak dimer interface is the locus of pathogenic mutations such as D18G, V20I, A25T, L111M, Y114C, and V122I that destabilize the tetramer and increase the rate of tetramer dissociation relative to that of WT TTR (15, 63–66).

The intensity of the tryptophan fluorescence emission maximum at 345 nm increases during the lag phase of agitation-induced aggregation (Figures 4A and 4D). The fluorescence intensity increases with increasing agitation time, indicating growth in the population of a perturbed tetramer with enhanced fluorescence emission. TTR contains two tryptophan residues, W41 and W79. At neutral pH, the intrinsic fluorescence of W79 in both the TTR tetramer and engineered monomeric M-TTR is quenched and the fluorescence spectrum is dominated by emission from W41 (15, 16, 49). The large increase in fluorescence emission intensity that we observe upon agitation of WT TTR most likely arises from dynamic changes that relieve quenching of the W79 fluorescence. The W79 side chain packs against residues in the AG and GH regions, where the intensity changes in HSQC cross peaks indicate enhanced dynamics following agitation. Thus, both the NMR and fluorescence measurements support a model in which agitation dynamically destabilizes the weak dimer interface.

The HSQC cross peaks of several residues lose intensity significantly more rapidly than average, likely due to exchange between the NMR-visible ground state and a weakly populated tetrameric excited state with altered conformation. The rate of intensity loss is greater than average for residues L12-V14 near the N-terminus of strand A, residues E54 and L55 in neighboring strand D, and residues in the DE loop that contact side chains in the N-terminal region of strand A. The perturbations also appear to propagate through side chain contacts to G47 and T49 on strand C and to the FG loop. Above average intensity loss is also observed for T123 and N124 at the C-terminus of the H strand and for the neighboring R104 on strand G. All of the above residues form an extensive cluster connected by backbone hydrogen bonds or by side chain contacts. The only “isolated” residue with greater than average broadening is F87, whose side chain projects across the strong dimer interface and docks into the neighboring protomer.

The dynamic perturbations observed in the NMR and fluorescence spectra of WT TTR could arise from shear forces acting directly on the tetramer or from incorporation of one or more agitation-damaged monomers into the reassembled tetramer. The second scenario seems more probable given that disruption of the tetramer interfaces leads to destabilization of the monomer and enhances local conformational fluctuations (33) that likely make the protein more susceptible to the effects of agitation. Except for the residues in strand C, all of the residues that undergo faster than average intensity loss upon agitation experience exchange broadening in spectra of the monomeric F87E TTR variant. R_2_ relaxation dispersion measurements show that broadening arises from conformational exchange between the F87E ground state and a weakly populated, amyloidogenic excited state in which the hydrogen bonding network of the DAGH β-sheet is perturbed (33). The conformational fluctuations, which are enhanced at acidic pH where amyloid formation is favored, transiently expose hydrophobic surfaces and predispose TTR towards local unfolding and progression into the aggregation pathway. The observation that the same set of residues experience exchange broadening in the monomer and loss of cross peak intensity following prolonged agitation of the tetramer suggests similar molecular mechanisms for structural remodeling and aggregation under acidic conditions or mechanical shear stress.

Several factors suggest that our observation of agitation-induced aggregation of WT human TTR at neutral pH has physiological relevance. Firstly, aggregation of the WT tetramer under agitation occurs on a physiologically relevant time scale and at concentrations within the typical range found in adult blood plasma (∼7-29 μM protomer concentration, half-life ∼48 hours) (39, 67–69). Plasma TTR concentrations decrease with age and low levels of TTR are a risk factor for cardiac amyloidosis; indeed, TTR levels less than ∼10 μM are associated with high risk or incidence of heart failure (69–72). At such low concentrations, the midpoint time for aggregation is less than 24 hours, well within the plasma half-life of TTR. The TTR concentration in cerebrospinal fluid is much lower (0.2 −1.6 μM), making it even more susceptible to aggregation induced by mechanical shear.

Normal vascular blood flow is laminar but is close to the onset of turbulence (28, 73, 74). Physiological shear forces associated with blood flow are too small to fully unfold small globular proteins (75, 76). However under conditions of turbulent flow, fluid motion becomes chaotic, generating high velocity gradients and intense shear and extensional forces that can induce transient exposure of hydrophobic residues, local unfolding, and protein aggregation (55, 56, 76, 77). Turbulent blood flow occurs at arterial branch points, in stenotic aortic valves and arteries, and in arteries that are partially obstructed by atherosclerotic plaques, and is exacerbated by the pulsatile nature of blood flow (28, 73, 74, 78–81) (79, 81–84). TTR cardiac amyloidosis in elderly patients is frequently associated with severe arterial and aortic valve stenosis (AS) (85–87) and it has been postulated that the increased mechanical stress associated with AS may initiate TTR amyloid formation and tissue infiltration (88, 89). Disturbed and turbulent flow also occurs in the cerebrospinal fluid (90), suggesting a potential role in TTR aggregation.

## Acknowledgements

This research was funded by grants DK124211 (PEW) and GM131693 (HJD) from the National Institutes of Health and by the Branco Weiss Fellowship – Science in Society (IR). We thank Ashok Deniz and Daniel Scholl for access to and support with light microscopy; Andrew Ward, Mengyu Wu and James Ferguson for negative stain EM imaging, Maria Martinez-Yamout, Xun Sun, James Ferguson, and Ben Leach for helpful discussions, and Gerard Kroon for support with the NMR spectroscopy.

