## Supplementary Material for "Aggregation of Transthyretin by Fluid Agitation"

### **Additional Experimental Details**

Protein Expression and Purification

Recombinant TTR constructs were overexpressed in *Escherichia coli* (BL21-DE3) in rich (LB broth) or $^{15}$N-enriched (1 g/l $^{15}NH_4Cl$) minimal (M9) medium and purified by two step Ni-affinity purification with enzymatic cleavage (TEV site) of an N-terminal $His_6$-tag. After growing cells to an O.D. of 0.8, protein over-expression was induced by addition of 1 mM IPTG, and cultures were harvested after 3-5 h continued shaking at 37°C (150 rpm). Cell pellets after centrifugation were resuspended in lysis buffer (20 mM Tris, pH 8.0, 1 M NaCl, 1 mM EDTA, 1 mM TCEP, 1 EDTA-free proteinase inhibitor cocktail tablet (cOmplete$^{TM}$, Roche, Switzerland), and frozen (-80°C ) until purification. For purification cell suspensions were thawed and lysed by sonication on ice. Lysates were clarified by centrifugation (30 min at 17k x g), and manual filtration (0.22 µM). Ni-affinity chromatography was performed using 5 ml (column volume after draining) cOmplete$^{TM}$ resin (Roche, Switzerland) in a gravity flow protocol. $His_6$-tev-TTR was eluted in a single step (250 mM imidazole). His-tag removal was performed at a ratio of ~1:100 (TEV protease:protein) after addition of fresh 0.5 mM TCEP and during overnight dialysis at room temperature to 20 mM Tris, pH 8.0, 100 mM NaCl. Without incorporation of at least one additional amino acid at position +1 of the TEV cleavage site, enzymatic cleavage was very inefficient. An additional Gly at position -1 (N-terminal of TTR) proved necessary and sufficient for complete enzymatic cleavage. After cleavage, the solution was re-passed over the Ni-affinity column, with the addition of 1M NaCl to ensure that all of the cleaved TTR would be collected in the flow-through. Protein identity and purity was confirmed by SDS-PAGE, HPLC, and MALDI mass spectrometry. By NMR we found that using high levels of reducing agent (1 mM TCEP) and addition of 0.5 mM EDTA during cell lysis and both Ni-column runs was critical to prevent TTR oxidation, otherwise a minor secondary peak (~10%-25%) was seen in HPLC.

Native electrophoretic mobility shift assay (EMSA)

For the EMSA screen, independent TTR samples were agitated under standard conditions (1200 rpm, agitation buffer) at $c_0$=5 µM (500 µl). A reference sample was kept on the bench without agitation. After 90 h, samples were centrifuged (30 min at 17k x g). Native gel loading dye was added to the soluble supernatant (final ~10% glycerol, <0.01 % bromophenol blue dye) and 15 µl samples were loaded onto an agarose gel (1% w/v in 25 mM Tris, 192 mM glycine, pH 8.3). EMSA was performed at 80 V for 40 min, followed by staining with Coomassie solution (10% methanol, 50% acetic acid, 0.01% Coomassie blue) and de-staining in deionized water.

Supplementary Tables

| $c_0$ [μM] | $k$ (ThT) [1 /h] | $I_p$ (*ThT*) | $I_p$ /$c_0$ (ThT) [1/μM] | $T_m$ (ThT) [h] | $R_{adj}$ (ThT) |
|---|---|---|---|---|---|
| WT TTR (TETRAMER) | | | | | |
| **5** | **0.2 ± 0.1** | **32 ± 3** | **6.4** | **20 ± 2** | **0.94** |
| **10** | **0.2 ± 0.1** | **53 ± 5** | **5.3** | **23 ± 1** | **0.95** |
| **20** | **0.022 ± 0.005** | **125 ± 9** | **6.3** | **145 ± 8** | **0.98** |
| 50 | 0.010 ± 0.004 | 318 ± 44 | 6.4 | 290 ± 30 | 0.96 |
| 100 | 0.012 ± 0.004 | 381 ± 41 | 3.8 | 290 ± 20 | 0.98 |
| F87E TTR (MONOMER) | | | | | |
| **5** | **0.12 ± 0.05** | **24 ± 3** | **4.8** | **23 ± 2** | **0.98** |
| **10** | **0.10 ± 0.02** | **70 ± 3** | **7.0** | **31 ± 2** | **0.99** |
| **20** | **0.08 ± 0.01** | **138 ± 3** | **6.9** | **51 ± 1** | **0.99** |

Table S1. Sigmoidal fit parameters of the $c_0$-dependent agitation screen for WT tetrameric TTR and F87E monomeric TTR, as observed by ThT fluorescence (note that due to intensity normalization, Ip is dimensionless); rows marked bold are fits with high fit quality that were analyzed in the main text; errors shown at 90% fit confidence interval.

| $c_0$ [μM] | $k_W$ ($I_{w,sol}$) [1/h] | $I_{W,0}$ / $10^5$ ($I_{w,sol}$) [cts] | $T_{m(W)}$ ($I_{w,sol}$) [h] | $R_{adjsq}$ |
|---|---|---|---|---|
| 0.5 | n.c. | n.c. | n.c. | |
| 1 | n.c. | n.c. | n.c. | |
| **5** | **0.14 ± 0.06** | **7 ± 1** | **21 ± 6** | **0.99** |
| 10 | n.c. | n.c. | n.c. | |
| **50** | **0.02 ± 0.02** | **8 ± 2** | **160 ± 50** | **0.96** |
| 100 | n.c. | 8 ± 2 | 170 ± 60 | 0.87 |

Table S2. Sigmoidal fit parameters of the $c_0$ dependent soluble Trp fluorescence screen with WT TTR; rows marked bold are fits for the data series shown in the main text; non-converged (n.c.) fit parameters were omitted (95% fit parameter error > 100% of fit parameter value). $I_0$ is shown as fitted and not back-calculated by dilution factor.

Table S3. Agitation assay for NMR analysis. Total $A_{330nm}$ is a direct measurement by nanodrop;

| time (days) | total $A_{330nm}$ | total $A_{600nm}$ | ThT fluorescence gain | average NMR intensity R(t) | average normalized NMR intensity $R_n(t)$ | supernatant Trp intensity ratio $I/I_{ref}$ |
|---|---|---|---|---|---|---|
| 0 | 0.014 | 0.10 ± 0.02 | 1.3 ± 0.2 | 1 (def) | 1.0 | 1.0 (def) |
| 4* | 0.185 | 0.23 ± 0.02 | 2.5 ± 0.3 | 0.91 | 0.90 | 2.7 |
| 6 | 0.093 | 0.15 ± 0.01 | 2.2 ± 0.3 | n.d. | n.d. | 1.7 |
| 10 | 0.221 | 0.21 ± 0.02 | 4.0 ± 0.8 | 0.92 | 0.87 | 1.5 |
| 14 | 0.272 | 0.23 ± 0.02 | 4.6 ± 1.0 | 0.89 | 0.86 | 2.4 |
| 16 | 0.361 | 0.21 ± 0.02 | 4.7 ± 0.6 | 0.86 | 0.85 | 2.1 |
| 21 | 0.441 | 0.30 ± 0.04 | 6.1 ± 1.0 | 0.90 | 0.82 | 2.7 |

total $A_{600nm}$ is from the plate reader, back-calculated from a 1:10 dilution; ThT gain is from the plate reader, a 1:10 dilution, with two dilutions per sample. The asterisk denotes a possible anomalous concentration in the 4-day sample, where $A_{330nm}$, $A_{600nm}$ and ThT fluorescence are all higher than might be expected from the results for the remaining members of the series. The native-gel electrophoresis performed on the NMR samples (Figure S6) provides evidence for such an anomalously high concentration.

Supplementary Figures

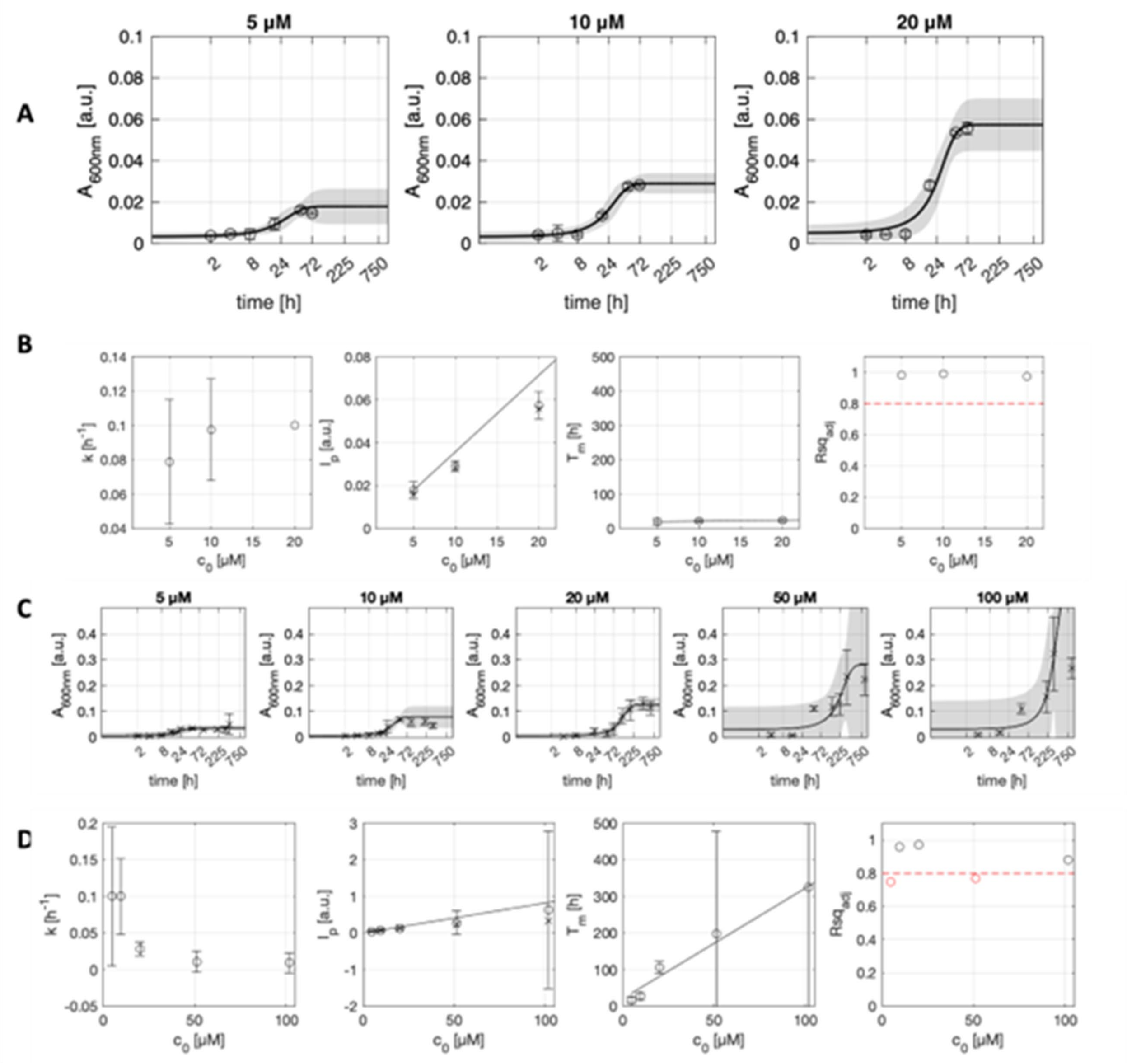


Figure S1. Time course and concentration dependence of TTR aggregation by acid (A, B) and agitation (C, D). (A) Acid aggregation data and fits; the $A_{600nm}$ values are not normalized; grey shaded areas are 75% confidence intervals of the fit; (B) fit parameters vs $c_0$; The $I_p$ panel shows the fitted parameter $I_p$ (o), as well as the actual experimental $A_{600nm,max}$ (x); solid lines where shown are linear regressions of the parameters versus $c_0$, where only parameters were considered where the global fit quality was sufficiently high ($Rsq_{adj}$>0.8, red dashed line). (C) Aggregation by agitation data and fits; the $A_{600nm}$ values are not normalized; grey shaded areas are 75% confidence intervals of the fit; (D) fit parameters vs $c_0$; The $I_p$ panel shows the fitted parameter $I_p$ (o), as well as the actual experimental $A_{600nm,max}$ (x); solid lines where shown are linear regressions of the parameters versus $c_0$, where only parameters were considered where the global fit quality was sufficiently high ($Rsq_{adj}$>0.8, red dashed line).

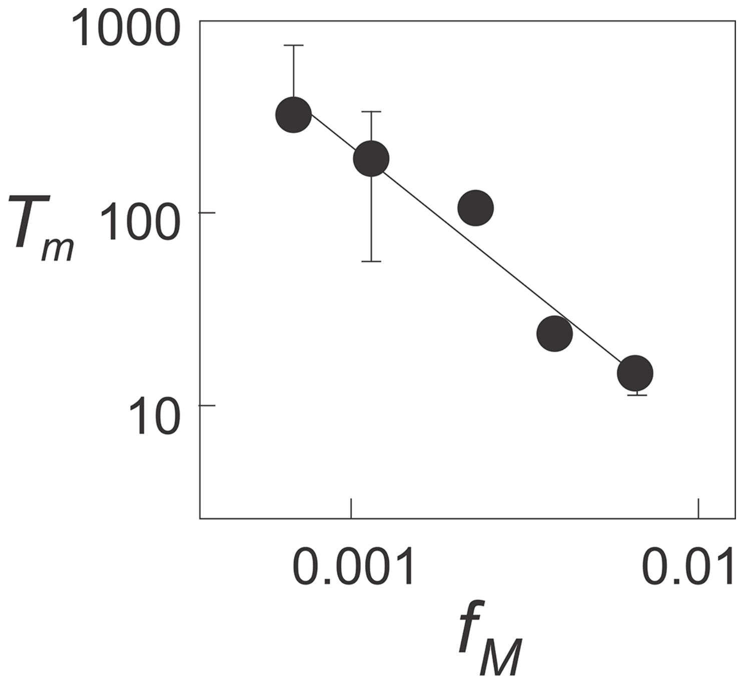


Figure S2. Plot of log of the lag time $T_m$ versus the fraction of monomer $f_M$.

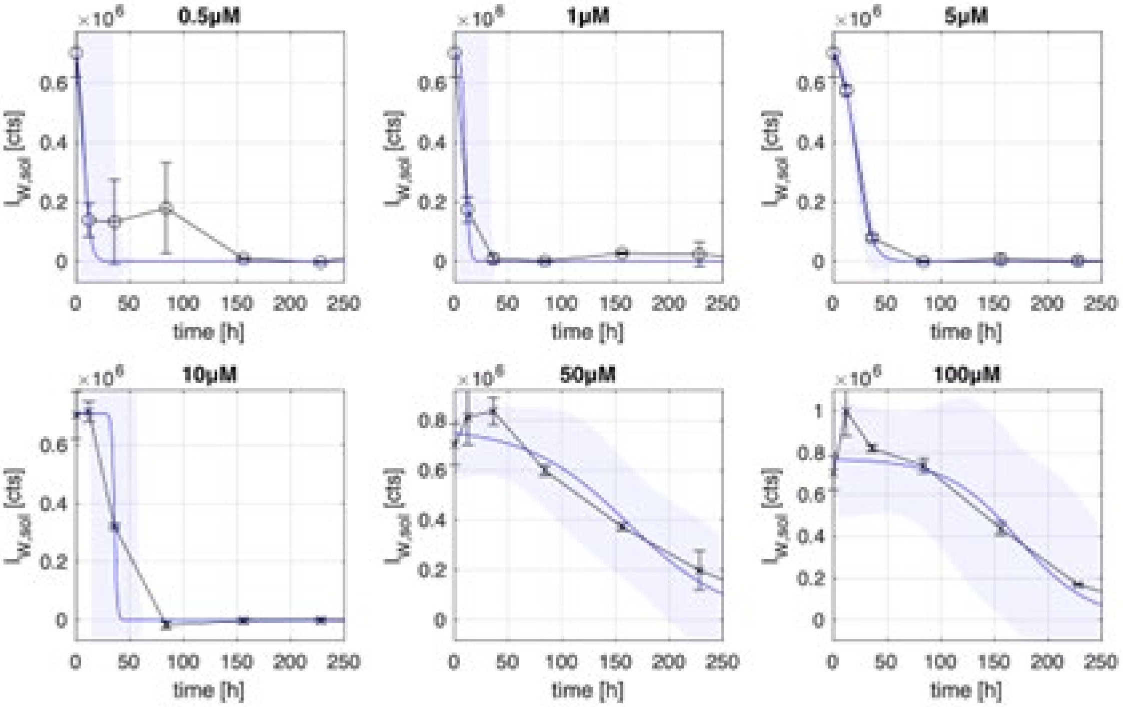


Figure S3. Full time courses (black) for soluble Trp fluorescence detected after agitation-induced aggregation of WT TTR at different $c_0$, with the sigmoidal fit according to Equation 2 (blue) and 90% confidence interval (shaded area).

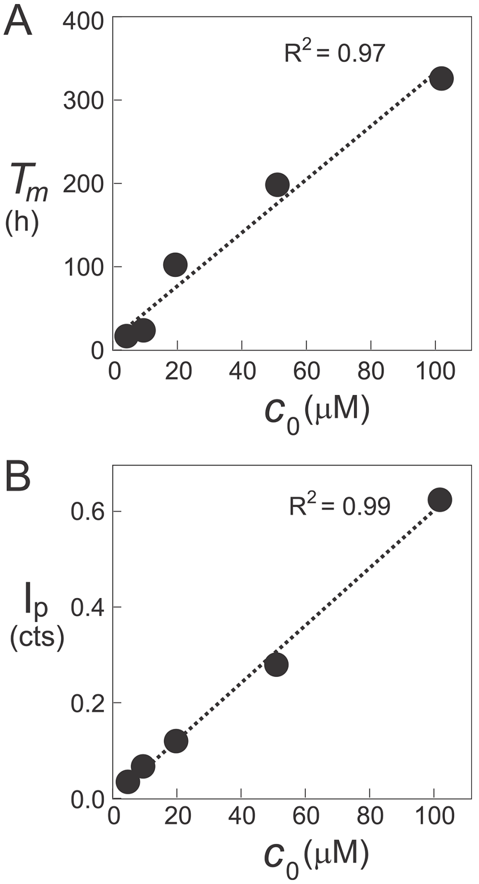


Figure S4. Concentration dependence of A. aggregation lag time $T_m$ and B. plateau intensity $I_p$ ($A_{600nm}$) for the samples used for NMR experiments.

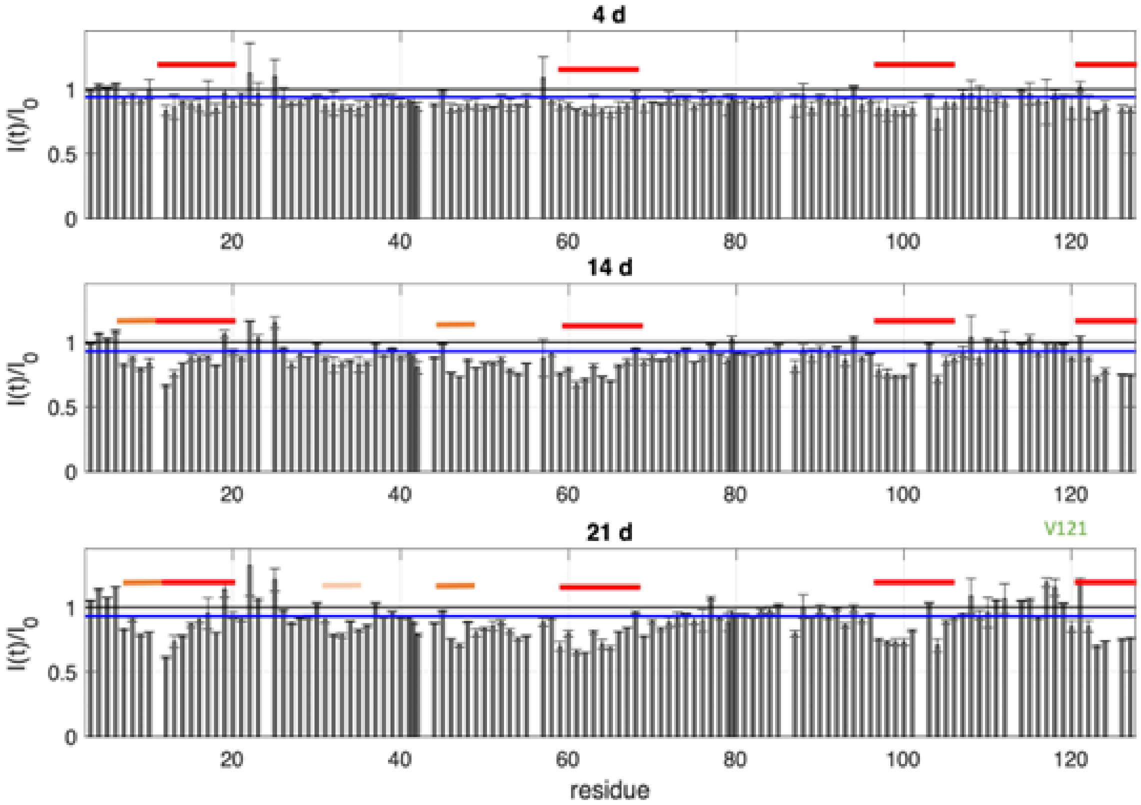


Figure S5. Unnormalized HSQC peak intensity ratios R(t) for three time points as indicated above; bars highlight ranges of substantial peak intensity loss (red: already weaker than average after 4 days, dark orange: weaker after 14 days, light orange: weaker after 21 days.

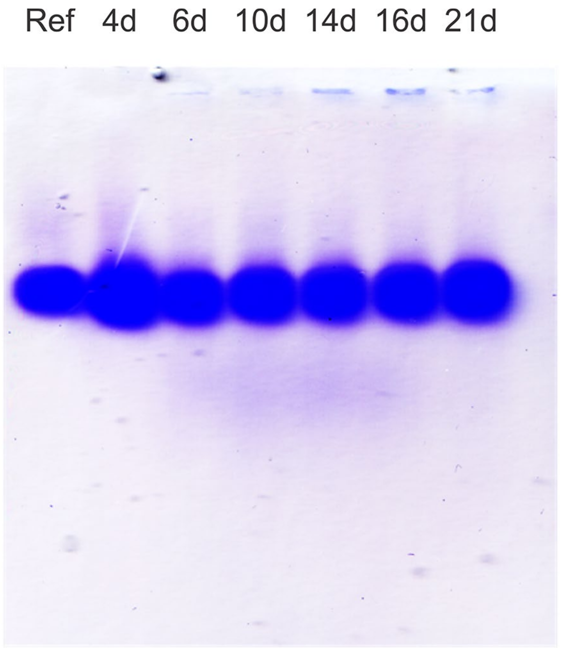


Figure S6. Native PAGE of NMR supernatant samples before agitation (Ref) and after 4, 5, 10, 14, 16, and 21 days of agitation. Note increased intensity of the 4 day sample, a possible indication that its concentration is anomalously high.
